# Suppressor analysis uncovers that MAPs and microtubule dynamics balance with the Cut7/Kinesin-5 motor for mitotic spindle assembly in *Schizosaccharomyces pombe*

**DOI:** 10.1101/380493

**Authors:** Masashi Yukawa, Yusuke Yamada, Takashi Toda

**Author notes:** **The corresponding author’s name, office mailing address including street name and number, and email address.** Takashi Toda: Hiroshima Research Center for Healthy Aging and Laboratory of Molecular and Chemical Cell Biology, Department of Molecular Biotechnology, Graduate School of Advanced Sciences of Matter, Hiroshima University, Hiroshima 1-3-1 Kagamiyama, Higashi-Hiroshima, 739-8530, JAPAN, Masashi Yukawa: Hiroshima Research Center for Healthy Aging and Laboratory of Molecular and Chemical Cell Biology, Department of Molecular Biotechnology, Graduate School of Advanced Sciences of Matter, Hiroshima University, Hiroshima 1-3-1 Kagamiyama, Higashi-Hiroshima, 739-8530, JAPAN.

## Abstract

The Kinesin-5 motor Cut7 in *Schizosaccharomyces pombe* plays essential roles in spindle pole separation, leading to the assembly of bipolar spindle. In many organisms, simultaneous inactivation of Kinesin-14s neutralizes Kinesin-5 deficiency. To uncover the molecular network that counteracts Kinesin-5, we have conducted a genetic screening for suppressors that rescue the *cut7-22* temperature sensitive mutation, and identified 10 loci. Next generation sequencing analysis reveals that causative mutations are mapped in genes encoding α-, β-tubulins and the microtubule plus-end tracking protein Mal3/EB1, in addition to the components of the Pkl1/Kinesin-14 complex. Moreover, the deletion of various genes required for microtubule nucleation/polymerization also suppresses the *cut7* mutant. Intriguingly, Klp2/Kinesin-14 levels on the spindles are significantly increased in *cut7* mutants, whereas these increases are negated by suppressors, which may explain the suppression by these mutations/deletions. Consistent with this notion, mild overproduction of Klp2 confers temperature sensitivity. Surprisingly, treatment with a microtubule-destabilizing drug not only suppresses *cut7* temperature sensitivity but also rescues the lethality resulting from the deletion of *cut7,* though a single *klp2* deletion per se cannot compensate for the loss of Cut7. We propose that microtubule assembly and/or dynamics antagonize Cut7 functions, and that the orchestration between these two factors is crucial for bipolar spindle assembly.

## INTRODUCTION

In eukaryotic cells, the assembly of mitotic spindles is a crucial step in accurate chromosome segregation. The mitotic spindle, a dynamic ensemble of microtubules, microtubule-associated proteins (MAPs) and motor proteins, ensures the proceeding of mitosis by aligning sister chromatids in the equatorial region and segregating them towards opposite poles (Mitchison and Salmon 2001; Woodruff *et al.* 2014). In order to assemble bipolar mitotic spindles, collaborative forces exerted by multiple kinesin motors, collectively called mitotic kinesins, are necessary (Yount *et al.* 2015).

The plus-end directed Kinesin-5 is essential for mitosis and, therefore, influences viability in many eukaryotes from fungi to human beings (budding yeast Cin8 and Kip1, fission yeast Cut7, *Aspergillus* BimC, *Drosophila* Klp61F, *Xenopus* Eg5 and human Kif11) (Enos and Morris 1990; Hagan and Yanagida 1990; Le Guellec *et al.* 1991; Heck *et al.* 1993; Blangy *et al.* 1995). The primary role of Kinesin-5 motors has been shown to be the establishment of spindle bipolarity by driving spindle pole separation. They form homotetramers that crosslink and subsequently slide apart antiparallel microtubules, thereby generating an outward pushing force to separate the two poles (Kashina *et al.* 1996; Kapitein *et al.* 2005). The sole member of the fission yeast Kinesin-5 family, Cut7, is indispensable for mitotic progression. Temperature-sensitive mutations in *cut7* display mitotic arrest with monopolar spindles at the restrictive temperature (Hagan and Yanagida 1990; Hagan and Yanagida 1992). This phenotypic consequence is identical to what is observed in other organisms, including human cells when Kinesin-5 activity is inhibited (Sawin *et al.* 1992; Kapoor *et al.* 2000). Because its function is essential to cell proliferation, Kinesin-5 molecules have been targeted for the development of anticancer drugs (Wacker *et al.* 2012; Ma *et al.* 2014; Dumas *et al.* 2016). However, at present, our knowledge on the physiology and regulation of Kinesin-5 motors is still too limited to draw a complete picture of their functions in vivo. It is therefore crucial to obtain a comprehensive understanding of how Kinesin-5 molecules establish spindle bipolarity, and which molecules regulate this kinesin.

Kinesin-5 motors are generally localized along spindle microtubules during mitosis. However, rather than having a uniform localization pattern on the mitotic spindle, the Kinesin-5 motors are enriched in two regions: at the medial midzone which corresponds to the microtubule plus ends, and near the centrosome which is proximal to the microtubule minus ends (Yount *et al.* 2015). Interestingly, recent work has shown that fungal Kinesin-5 motors are bi-directional: they can move processively towards both plus- and minus-end direction on the microtubules under various conditions (Gerson-Gurwitz *et al.* 2011; Roostalu *et al.* 2011; Edamatsu 2014; Britto *et al.* 2016). It has been reported that individual molecules of budding yeast Cin8 and fission yeast Cut7 initially move towards the minus end, and when concentrated/crowed in clusters on the microtubule, they switch motility from a minus end- to plus-end-directed manner, thereby generating robust outward force (Britto *et al.* 2016; Shapira *et al.* 2017). This bi-directionality and motility switch may account for the biased localizations of Kinesin-5 on the spindle, though presently, its physiological relevance and significance remain to be established.

Mitotic spindle assembly, i.e. the formation of a microtubule bipolar array, requires another Kinesin family motor, Kinesin-14, which exhibits a minus-end directed motility (She and Yang 2017. In fission yeast, two Kinesin-14s, Pkl1 and Klp2, counteract Cut7-dependent spindle pole separation (Pidoux *et al.* 1996; Troxell *et al.* 2001; Furuta *et al.* 2008; Braun *et al.* 2009). Pkl1 is localized predominantly to the mitotic spindle pole body (SPB, the fungal equivalent of the animal centrosome), where it forms a ternary complex with Msd1 and Wdr8 (referred to as the MWP complex) (Toya *et al.* 2007; Ikebe *et al.* 2011; Syrovatkina and Tran 2015; Yukawa *et al.* 2015). During early mitosis when duplicated SPBs initially separate, this anchorage generates inward force at the SPB against Cut7-mediated outward force (Yukawa *et al.* 2018). Klp2, on the other hand, is localized to the spindles in a punctate manner and crosslinks microtubule bundles using the N-terminal-located second microtubule binding domain (Braun *et al.* 2009), by which it generates inward forces at the SPBs. Accordingly, the deletion of either *pkl1* or *klp2* suppresses the temperature sensitivity caused by the *cut7* mutations (Pidoux *et al.* 1996; Troxell *et al.* 2001; Rodriguez *et al.* 2008). It is noteworthy that although Pkl1 and Klp2 collaboratively antagonize Cut7, their contributions are not identical; while the *pkl1* deletion *(pkl1Δ)* can completely bypass the requirement of Cut7’s essentiality, *klp2Δ* is incapable of rescuing *cut7Δ* (Yukawa *et al.* 2018).

In this study we sought to identify, in an unbiased manner, genes that antagonize the outward force generated by Cut7/Kinesin-5. To this end, we screened for extragenic suppressors that are capable of rescuing the temperature sensitivity of the hypomorphic *cut7* mutants. Consequently, we identified 6 suppressor genes. Subsequent analysis of these genes led us to the proposition that in addition to the MWP complex, microtubule stability and/or dynamics play a crucial role in antagonizing Cut7-mediated outward force. In fact, we have found that chemical perturbation of microtubule stability/dynamics is capable of suppressing the hypomorphic *cut7* mutant phenotype. Remarkably, we have further discovered that treatment with a microtubule destabilizing drug is sufficient to bypass the essential function of Cut7. We discuss how microtubule stability and/or dynamics keep inward and outward forces in balance, thereby assembling bipolar mitotic spindles.

## MATERIALS AND METHODS

### Strains, media, and genetic methods

Fission yeast strains used in this study are listed in Supplemental Table S1. Media, growth conditions, and manipulations were carried out as previously described (Moreno *et al.* 1991; Bähler *et al.* 1998; Sato *et al.* 2005). For most of the experiments, rich YE5S liquid media and agar plates were used. Wild-type strain (513; Supplemental Table S1), temperature-sensitive *cut7-22,* and *pkl1* deletion strains were provided by P. Nurse (The Francis Crick Institute, London, England, UK), I. Hagan (Cancer Research UK Manchester Institute, University of Manchester, Manchester, England, UK), and R. McIntosh (University of Colorado, Boulder, CO), respectively. Spot tests were performed by spotting 5–10 μl of cells at a concentration of 2 × 10^7^ cells/ml after 10-fold serial dilutions onto rich YE5S plates with or without a drug (TBZ). Some of the YE5S plates also contained Phloxine B, a vital dye that accumulates in dead cells and stains the colonies dark pink (Moreno *et al.* 1991). The plates were incubated at various temperatures from 27°C to 36°C as necessary.

### Preparation and manipulation of nucleic acids

Enzymes were used as recommended by the suppliers (New England Biolabs inc., Ipswich, MA and Takara Bio Inc., Shiga, Japan).

### Strain construction, gene disruption, and the N-terminal and C-terminal epitope tagging

A PCR-based gene-targeting method (Bähler *et al.* 1998; Sato *et al.* 2005) was used for complete gene disruption and GFP-tagging.

### Isolation of revertants from the *cut7* mutants

Approximately 2 × 10^7^ *cut7-22* cells/plate were spread on YE5S plates (>20 plates) and directly incubated at 36°C. After incubation for several days, revertant colonies were picked up, streaked on YES5 plates and incubated at 36°C. Revertants were backcrossed with a *cut7^+^-GFP-kanR* wild-type strain and individual segregants were spread on YE5S plates. If no Ts^-^ segregants appeared, these were assigned as intragenic suppressors, while if Ts^-^ segregants appeared, they were scored as extragenic suppressors. Following this, pair-wise crosses were performed among individual extragenic suppressors. If no Ts^-^ segregants appeared, these two suppressors are classified to belong to the same linkage group. In this way, in total 10 *skf* loci were assigned. Next, one representative allele from individual *cut7-22 skf* mutants were crossed with a strain containing deletion of *pkl1, msd1* or *wdr8.* If Ts^-^ segregants appeared, it is judged that this *skf* gene is not allelic to *pkl1, msd1* or *wdr8.* From this analysis, it is concluded that *skf1*, *skf2* and *skf3* are *pkl1*, *wdr8* and *msd1*, respectively, which was confirmed by nucleotide sequencing of these genes in corresponding *skf* mutants.

### Next generation sequencing and annotation of *skf* genes

At least 5 independent colonies of each *skf* mutant were mixed and cultured in the liquid YE5S. A parental *cut7-22* mutant was also cultured. Genomic DNA was purified according to a standard method (IIDA *et al.* 2014). Whole genome sequencing was performed by BGI Next Generation Sequencing (NGS) services (Shenzhen, China). Individual sequence data were analyzed using Mudi (Mutation discovery, http://naoii.nig.ac.jp/muditop.html) (IIDA *et al.* 2014).

### Fluorescence microscopy and time-lapse live cell imaging

Fluorescence microscopy images were obtained by using the DeltaVision microscope system (DeltaVision Elite; GE Healthcare, Chicago, IL) comprising a wide-field inverted epifluorescence microscope (IX71; Olympus, Tokyo, Japan) and a Plan Apochromat 60×, NA 1.42, oil immersion objective (PLAPoN 60×O; Olympus Tokyo, Japan). DeltaVision image acquisition software (softWoRx 6.5.2; GE Healthcare, Chicago, IL) equipped with a charge-coupled device camera (CoolSNAP HQ2; Photometrics, Tucson, AZ) was used. Live cells were imaged in a glass-bottomed culture dish (MatTek Corporation, Ashland, MA) coated with soybean lectin and incubated at 27°C for most of the strains and at either 27°C, 33°C or 36°C for the ts mutants. The latter two cultures were incubated in rich YE5S media until mid–log phase at 27°C and subsequently shifted to the restrictive temperature of either 33°C or 36°C before observation. To keep cultures at the proper temperature, a temperature-controlled chamber (Air Therm SMT; World Precision Instruments Inc., Sarasota, FL) was used. The sections of images were compressed into a 2D projection using the DeltaVision maximum intensity algorithm. Deconvolution was applied before the 2D projection. Images were taken as 14–16 sections along the z axis at 0.2-μm intervals; they were then deconvolved and merged into a single projection. Captured images were processed with Photoshop CS6 (version 13.0; Adobe, San Jose, CA).

### Quantification of fluorescent signal intensities

For quantification of signals intensities of fluorescent marker-tagged protein (e.g. GFP-Klp2) located on the spindle microtubules, 14–16 sections were taken along the z-axis at 0.2-μm intervals. After deconvolution, projection images of maximum intensity were obtained, and after subtracting background intensities, values of maximum fluorescence intensities were used for statistical data analysis.

### Statistical data analysis

We used the two-tailed unpaired Student’s t-test to evaluate the significance of differences in different strains. All the experiments were performed at least twice. Experiment sample numbers used for statistical testing were given in the corresponding figures and/or legends. We used this key for asterisk placeholders to indicate p-values in the figures: e.g., ****, P < 0.0001.

### Data Availability

Strains and plasmids are available upon request. The authors affirm that all data necessary for confirming the conclusions of the article are present within the article, figures, and tables.

## RESULTS

### Isolation of revertants from the *cut7-22* temperature sensitive mutants

In order to identify genes counteracting the Cut7/Kinesin-5 function, we isolated spontaneous mutants capable of suppressing the *cut7-22* temperature-sensitive (ts) growth defect at 36°C. This screening yielded in total 25 revertants. Subsequent genetic analysis, including backcrossing with a wild-type *cut7* strain, defined that 3 isolates are intragenic, while the remaining 22 revertants contain extragenic suppressors. Nucleotide sequencing of the *cut7* gene in the 3 intragenic suppressors showed that each revertant contains one additional point mutation within the *cut7 ORF* besides the original mutation (P1021S) (Olmsted *et al.* 2014; Yukawa *et al.* 2018); two mutations were located at the stalk domain of Cut7 (R433Q and L687F), while the third mutation resided at the tail domain (S1027L) (Supplemental Figure S1A).

Pair-wise crosses between each of the 22 strains carrying extragenic suppressors indicated that these mutations consist of 10 linkage groups, designated *skf1-skf10 (skf:* suppressor of kinesin five). To identify causative mutations within extragenic suppressors, we performed next generation sequencing and analyzed whole-genome sequence data using the Mudi (Mutation discovery), a web tool for identifying mutations based on bioinformatics approaches (Iida *et al.* 2014). This analysis successfully identified individual genes of *skf1-skf6* (Figure 1A and B and Supplemental Figure 1B). These 6 genes could be categorized into two functional groups: Group I *(skf1-skf3)* comprises of genes encoding the components of the MWP complex (Skf1/Pkl1, Skf2/Wdr8 and Skf3/Msd1) as we expected (Yukawa *et al.* 2015), while Group II *(skf4-skf6)* includes those encoding microtubule constituents such as β-tubulin (Skf4/Nda3) (Hiraoka *et al.* 1984), α2-tubulin (Skf5/Atb2) (Toda *et al.* 1984) and the microtubule plus-end tracking protein EB1 homolog (Skf6/Mal3) (Beinhauer *et al.* 1997; Carvalho *et al.* 2003; Asakawa *et al.* 2005). The mutation sites within α2- and β-tubulin molecules correspond to the highly conserved amino acids that are located at the outer surface of the α/β-tubulin heterodimer (Supplemental Figure 1C and D). Consistent with these assignments, each deletion of *skf1-skf6* genes suppressed the *cut7-22* ts growth defect at 36°C (Figure 1C).

**Figure 1.**
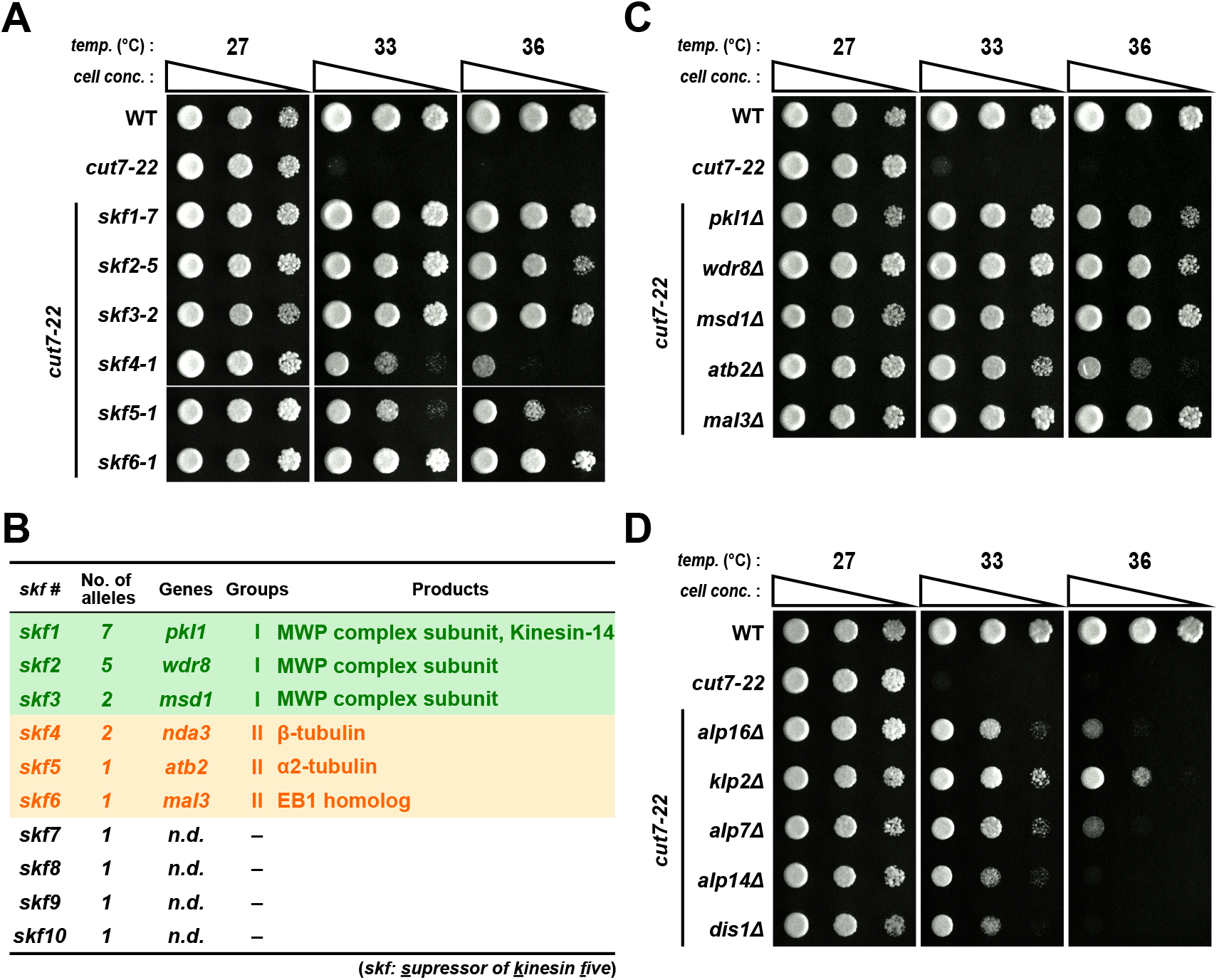
Identification of suppressor genes that rescue the *cut7-22* temperature sensitive mutant. **(A)** Spot test. Indicated strains were spotted onto rich YE5S agar plates and incubated at 27°C, 33°C or 36°C for 3 d. 10-fold serial dilutions were performed in each spot. *cell conc.,* cell concentration, *temp.,* temperature. **(B)** A summary table of *skf* genes. Group I includes genes encoding the MWP complex, while those in Group II encode either tubulins or the microtubule plus-end tracking protein Mal3/EB1. *n.d.:* not determined. **(C, D)** Spot tests. The same procedures were followed as in **(A)**.

Given that mutations/deletions of genes encoding tubulins or the EB1 homolog suppressed the growth defect caused by *cut7-22*, we next examined whether deletions of other genes involved in microtubule assembly and/or dynamics also display suppressing activity toward *cut7-22*. For this purpose, we examined gene deletions of *alp16* (encoding an ortholog of GCP6, a component of the γ-tubulin complex, γ-TuC) (Fujita *et al.* 2002; Anders *et al.* 2006; Masuda and Toda 2016), *alp7* (encoding an ortholog of the transforming acidic coiled-coil protein TACC) (Sato *et al.* 2004), and *alp14* and *dis1* (encoding XMAP215/TOG microtubule polymerases) (Nabeshima *et al.* 1995; Garcia *et al.* 2001; Al-Bassam *et al.* 2012; Hussmann *et al.* 2016; Matsuo *et al.* 2016), all of which play crucial roles in promoting proper mitotic spindle assembly.

Intriguingly, these deletions individually rescued the *cut7-22* ts mutants (Figure 1D). These results indicate that in addition to the impairment of the Pkl1 minus-end directed motor, the perturbation of spindle microtubule assembly including nucleation, polymerization and stabilization, also rescues the *cut7* mutant.

### Klp2/Kinesin-14 levels on the spindles determine temperature sensitivity of the *cut7* mutant

We previously showed that two Kinesin-14s, Pkl1 and Klp2, generate collaborative inward forces against the outward force exerted by Cut7/Kinesin-5; the deletion of either of these two genes suppresses the *cut7* ts mutants (Yukawa *et al.* 2018). Indeed, Group I suppressors include genes encoding the Pkl1-containing MWP complex (Figure 1B). Hence, one plausible scenario of rescuing the *cut7-22* mutation by Group II suppressors *(skf4-skf6)* would be the downregulation of Kinesin-14 function. To investigate this possibility, we measured the levels of Kinesin-14s, Pkl1 and Klp2, in various strains, at mitotic SPBs and on spindle microtubules, respectively. Interestingly, we found that Klp2 levels were substantially lessened by either *mal3Δ* or *alp16Δ,* whereas conversely, those in the *cut7-22* single mutant were increased (Figure 2A and B).

**Figure 2.**
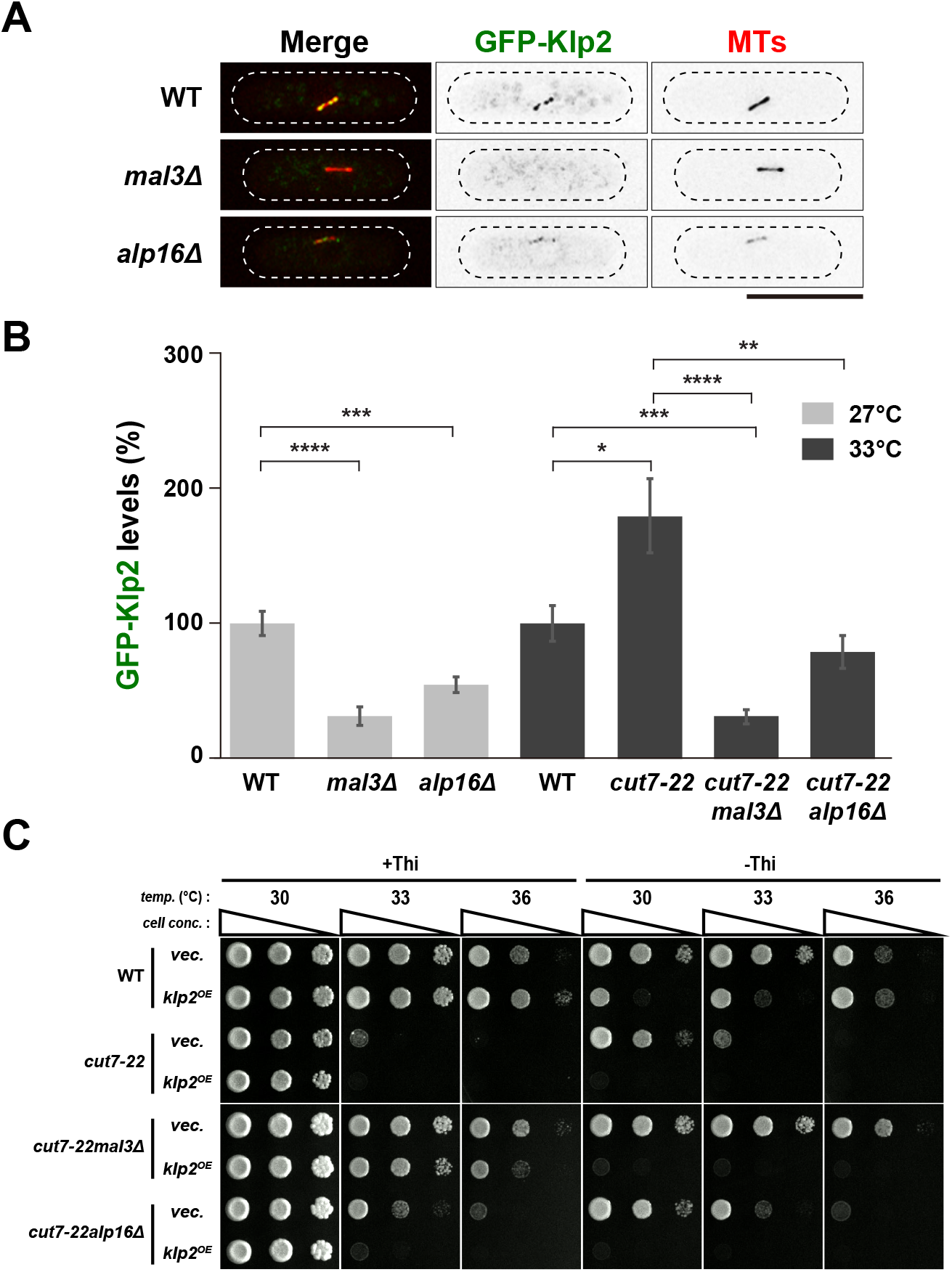
Klp2/Kinesin-14 levels on the spindles determine temperature sensitivity of the *cut7* mutant. **(A)** Representative images showing mitotic localization of GFP-Klp2 on the spindle microtubule are presented in indicated cells. All strains contain GFP-Klp2 and mCherry-Atb2 (a microtubule marker). Cells were incubated at 27°C. The cell peripheries are outlined with dotted lines. Scale bar, 10 μm. **(B)** Quantification of GFP-Klp2 levels on the spindle microtubule. Individual strains shown in **(A)** were grown at 27°C and a half of each culture was shifted to 33°C, while the remaining half was kept at 27°C. Fluorescence intensities were measured 2 h later. 33°C was used as the restrictive temperature, as fluorescence signals of GFP-Klp2 were quenched rapidly at 36°C, which made quantitative measurement of GFP-Klp2 signal intensities difficult. The total values of GFP-Klp2 fluorescence intensities were divided by the spindle length in each cell. The values obtained from wild-type cells incubated at 27°C and 33°C were set as 100%, and compared to those from other strains incubated at the same temperature. All p-values were obtained from the two-tailed unpaired Student’s t test. Data are presented as the means ± SE (≥16 cells). *, P < 0.05, **, P < 0.01, ***, P < 0.001, ****, P < 0.0001. (C) Spot test. Individual strains containing a plasmid carrying the *klp2+* gene under the thiamine-repressible *nmt1* promoter (Maundrell 1990) or an empty vector were spotted on minimal plates in the presence (+Thi, 10 μg/ml) or absence (-Thi) of thiamine, and incubated at various temperatures for 3 d. 10-fold serial dilutions were performed in each spot. *cell conc.,* cell concentration, *temp.,* temperature. Note that temperature sensitivity of *cut7-22mal3Δ* or *cut7-22alp16Δ* became exacerbated upon mild overproduction of *klp2+* (33°C or 36°C). Excessive overproduction of Klp2 (-Thi) was lethal in all the strains at any temperature (Yukawa *et al.* 2018).

Intriguingly, in the double mutants (*cut7-22mal3Δ* or *cut7-22alp16Δ*), Klp2 levels were significantly lower than those in *cut7-22* (Figure 2B). The reduction of Klp2 intensities on the spindle microtubules could not be attributed to the structural defects in the spindle microtubules, as midzone markers including Klp9/Kinesin-6 and Ase1/PRC1 (the anaphase B kinesin and the microtubule antiparallel crosslinker, respectively) (Loiodice *et al.* 2005; Yamashita *et al.* 2005; Fu *et al.* 2009) were properly localized to this region in *mal3Δ* cells (Supplemental Figure S2A and B). It is of note that the requirement of Mal3 for the microtubule localization of Klp2 has been previously reported (Mana-Capelli *et al.* 2012), which is in line with our current result. In contrast, Pkl1 levels at mitotic SPBs remained largely unaltered or even became higher in all the strains examined (Supplemental Figure S3A and B).

If the reduced levels of Klp2 on the spindle microtubules were responsible for the suppression of the ts phenotype of *cut7-22* by *mal3Δ* or *alp16Δ,* increased dosage of Klp2 in the double mutants might confer temperature sensitivity again. This was indeed the case. As shown in Figure 2C, mild overproduction of Klp2 conferred ts growth of *cut7-22mal3Δ* or *cut7-22alp16Δ* double mutants (Figure 2C). These observations imply that the reduced localization of Klp2 at the spindle microtubules, which results from mutations in *mal3* or *alp16*, is the main, if not the sole, reason for suppression of the *cut7* mutants. We envision that in the *cut7* mutants, the increased accumulation of Klp2 on the spindle microtubules contributes to the generation of excessive inward force.

### Intensities of spindle microtubules are increased in the *cut7-22* mutant but reduced in the absence of Mal3/EB1 or Alp6/GCP6

In the *cut7-22* mutant, more Klp2 proteins accumulate on the spindles. Moreover, we previously showed that this mutant displays hyper-resistance to a microtubule-destabilizing drug, thiabendazole (TBZ) (Yukawa *et al.* 2015). These findings raised the possibility that microtubule numbers are increased, by which spindle microtubules become more stable in the *cut7* mutant than in wild-type cells, leading to Klp2 accumulation and TBZ resistance. To interrogate this point, we directly measured fluorescence signal intensities of mCherry-Atb2 on the spindle microtubule. Intriguingly, the levels of preanaphase spindle microtubules (< 3 μm) were higher in the *cut7* mutant cells than those in wild-type cells at both the permissive and restrictive temperature, 27°C and 36°C respectively (Figure 3A-C and Supplemental Figure S4A and B).

**Figure 3.**
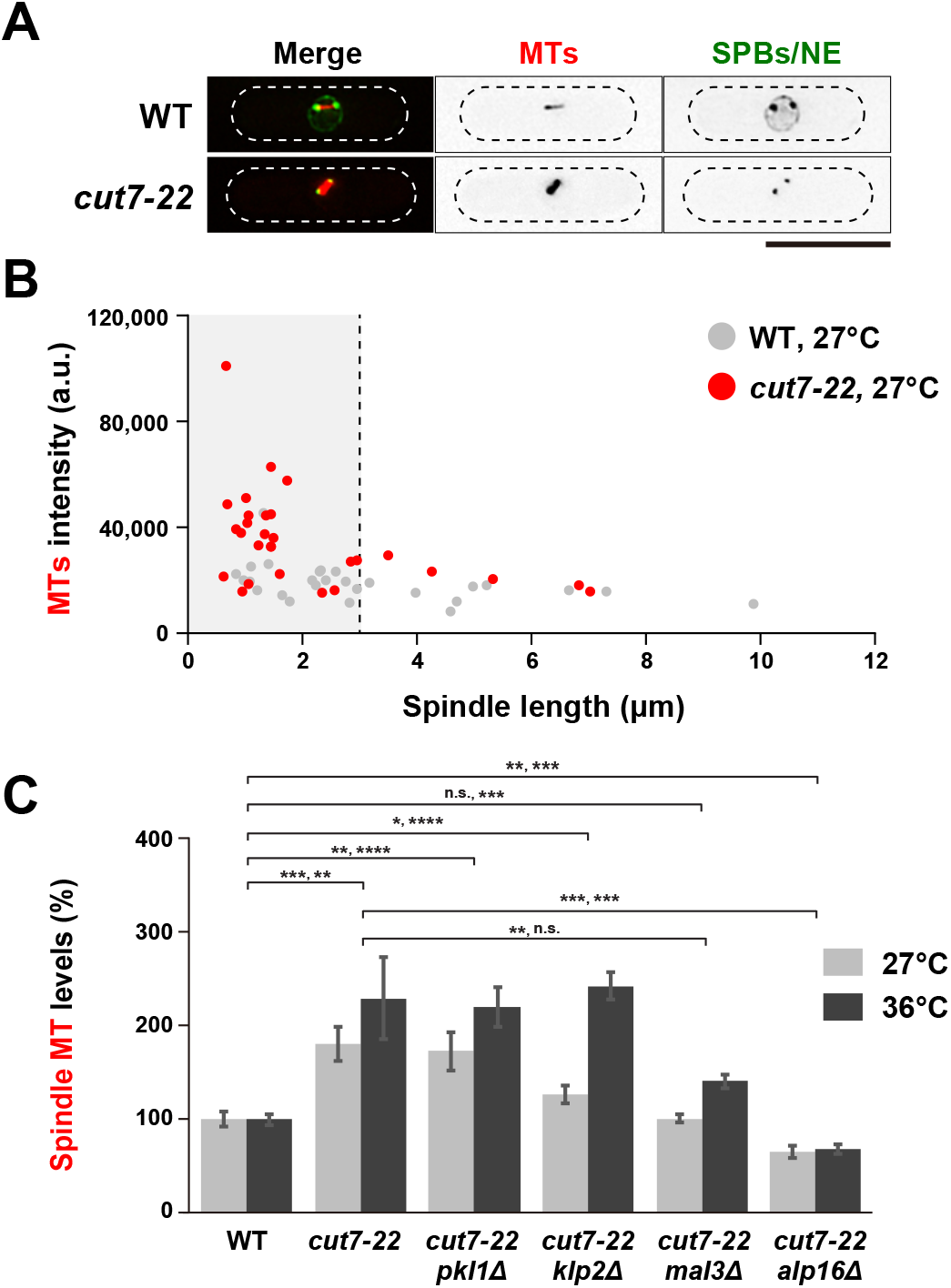
Intensities of the spindle microtubule are increased or conversely reduced in *cut7-22* or mutations that suppress *cut7-22*, respectively. **(A)** Representative images showing preanaphase spindle microtubules. While wild-type cells contain mCherry-Atb2 (MTs), GFP-Alp4 (SPBs) and Cut11-GFP (a marker for the nuclear membrane and mitotic SPBs) (West *et al.* 1998), while *cut7-22* cells contain mCherry-Atb2 and GFP-Alp4. Both strains were incubated at 27°C and preanaphase cells (spindle length is < 3 μm) were imaged. Scale bar, 10 μm. **(B)** Distribution of spindle microtubule intensities in wild-type (gray circles) and *cut7-22* cells (red circles). Spindle microtubule levels of individual mitotic cells shown in **(A)** were plotted against the spindle length. A vertical dotted line represents the spindle length (3 μm) at metaphase. **(C)** Quantification of the spindle microtubule. Fluorescence intensities of mCherry-Atb2 were measured in indicated strains that were incubated at either 27°C or 36°C. Most (>80%) of *cut7-22* mutant cells arrested with monopolar spindles; therefore, spindle intensities of mutant cells displaying short spindles were measured. The values of wild-type cells incubated at 27°C or 36°C were set as 100%, and compared to those from other strains incubated at 27°C or 36°C, respectively, for 2 h. All p-values were obtained from the two-tailed unpaired Student’s t test. Data are presented as the means ± SE (≥11 cells). *, P < 0.05, **, P < 0.01, ***, P < 0.001, ****, P < 0.0001. n.s., not significant.

We further found that the spindle intensities remained higher in the *cut7-22pkl1Δ* or *cut7-22klp2Δ* double mutant, but they were significantly reduced in either of *cut7-22alp16Δ* or *cut7-22mal3Δ* cells (Figure 3C). These results suggest that Klp2 accumulation on the spindles in the *cut7* mutant cells might be attributable to increased spindle levels and that its reduced accumulation in *cut7-22alp16Δ* or *cut7-22mal3Δ* leads to comprised inward force, resulting in the suppression of temperature sensitivity. It is also noted that the activity of spindle microtubule nucleation/assembly is potentiated when Cut7/Kinesin-5 function is compromised independently of Klp2. However, the levels of Alp4/GCP2, a core component of the γ-TuC (Vardy and Toda 2000) per se, were not increased in the *cut7* mutants (Supplemental Figure S4C and D). This result indicates that the potentiation of spindle assembly/nucleation in the *cut7* mutants, if this is the case, does not result from the accelerated recruitment of the microtubule nucleator γ-TuC to the mitotic SPBs.

### Drug-induced microtubule destabilization rescues the *cut7* temperature sensitive mutation with reduced Klp2 levels at the spindle microtubule

As the temperature sensitivity of *cut7-22* mutants is suppressed by the deletion of various genes that promote microtubule nucleation, assembly and organization, we presumed that simple microtubule destabilization by treatment with anti-microtubule drugs might also effectively rescue the growth defect of the *cut7* ts mutant cells. Therefore, we assessed the impact of TBZ treatment on *cut7* ts mutant strains. As shown in Figure 4A, growth properties of not only *cut7-22* but also all the other *cut7* ts alleles examined were ameliorated on TBZ-containing plates (Figure 4A).

**Figure 4.**
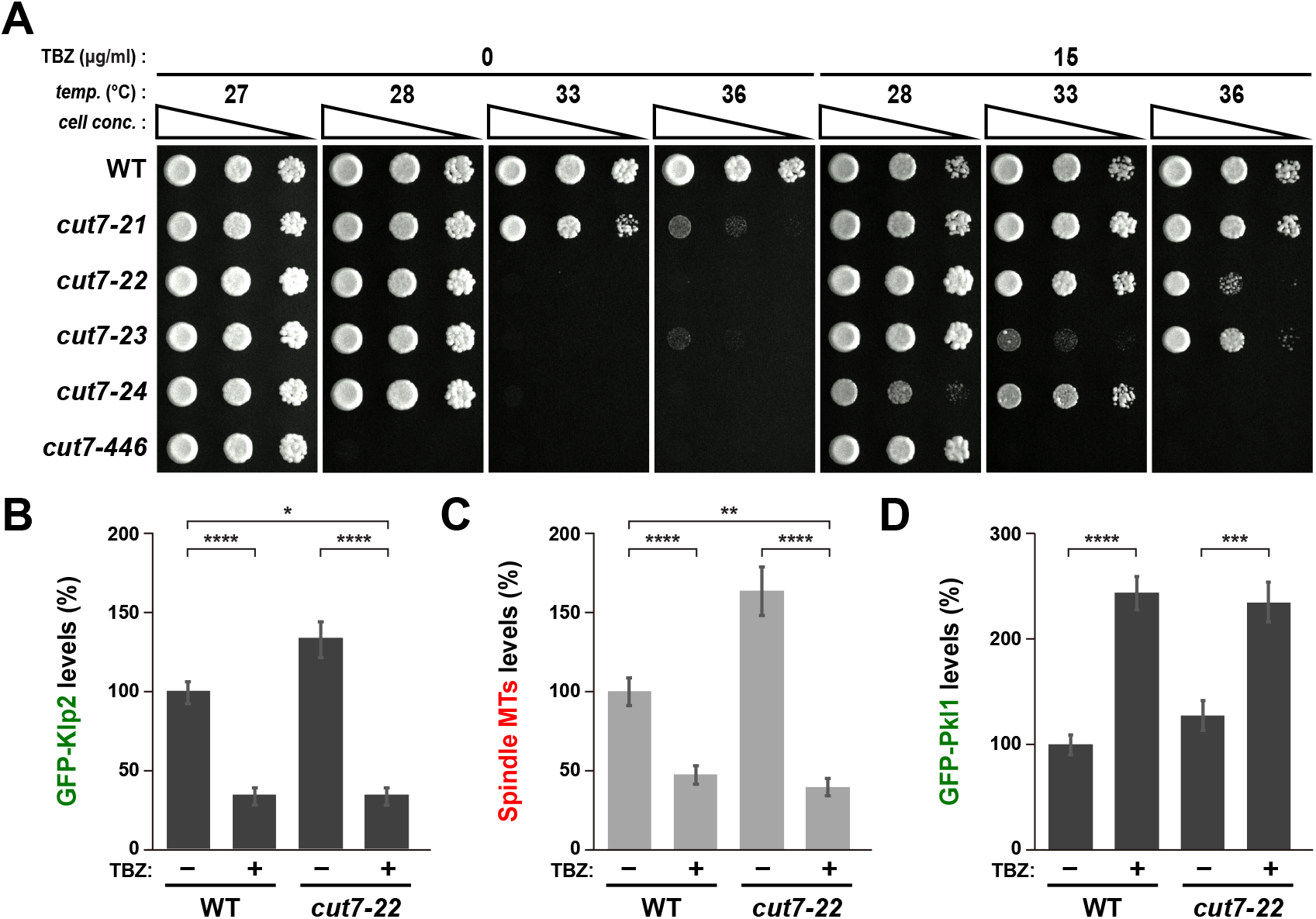
Treatment of a microtubule-destabilizing drug rescues various *cut7* temperature sensitive mutants with reduced levels of Klp2 at the mitotic spindle. **(A)** Spot test. Indicated strains were spotted onto rich YE5S agar plates in the absence or presence (15 μg/ml) of TBZ, and incubated at 27°C, 28°C, 33°C or 36°C for 3 d. 10-fold serial dilutions were performed in each spot. *cell conc.,* cell concentration, *temp.,* temperature. **(B-D)** Quantification. Fluorescence intensities of GFP-Klp2 on the spindle microtubule **(B)**, spindle microtubules **(C)** and GFP-Pkl1 at the mitotic SPB were measured in wild-type and *cut7-22* cells that were incubated at 27°C for 12-16 h in the absence or presence of 20μg/ml TBZ. All p-values were obtained from the two-tailed unpaired Student’s t test. Data are presented as the means ± SE (>10 cells). *, P < 0.05, **, P < 0.01, ***, P < 0.001, ****, P < 0.0001.

Given that the intensities of spindle-localizing Klp2 and those of the spindles themselves are proportionate and more importantly, that their levels correlate with whether *cut7-22* displays temperature sensitivity (higher) or not (lower) (see Figures 2B and 3C), we reasoned that TBZ treatment would decrease Klp2 levels on mitotic spindles. Indeed, we found that in the liquid culture, the Klp2 levels on the spindles as well as spindle microtubule intensities were reduced (Figure 4B and C). In clear contrast, the Pkl1 levels at mitotic SPBs were not decreased or even became higher with TBZ treatment (Figure 4D). This result indicates that microtubule destabilization by either mutations in genes encoding certain MAPs or a chemical reagent could rescue the *cut7* ts phenotype.

### Microtubule destabilization by a drug even bypasses the essentiality of Cut7/Kinesin-5 for viability

We then performed tetrad dissection of diploids heterozygous for *cut7* and *pkl1 (cut7^+^/Δ Δ/pkl1^+^),* and allowed spores to germinate on plates containing TBZ. Remarkably, this procedure rendered otherwise lethal *cut7Δ* haploid segregants viable; *cut7Δ* cells formed colonies, despite the fact that their size was much smaller than that of wild-type or *cut7Δpkl1Δ* doubly deleted segregants (Figure 5A) (Olmsted *et al.* 2014; Syrovatkina and Tran 2015; Yukawa *et al.* 2017). Continuous cell division of *cut7Δ* cells recovered from TBZ-containing plates upon germination still required the presence of this drug; no colonies were formed on plates in the absence of TBZ (Figure 5B). This indicates that the viability of *cut7Δ* cells is not ascribable to the emergence of suppressor mutations (e.g. *pkl1* or *msd1)* during the period of initial TBZ treatment. We found that very low concentrations of TBZ (2.5 μg/ml), in which no growth defects were apparent in wild-type cells, are sufficient to confer the viability of *cut7Δ* cells. At high concentrations of TBZ (>15 μg/ml), rescuing activity became compromised, as the growth of even wild-type cells was substantially inhibited under these conditions (Figure 5B). Therefore, a moderate disturbance of microtubule stability is crucial to bypass the requirement of Kinesin-5 function. Collectively, the impairment of microtubule stability and/or dynamics renders fission yeast cells viable in the absence of Kinesin-5.

**Figure 5.**
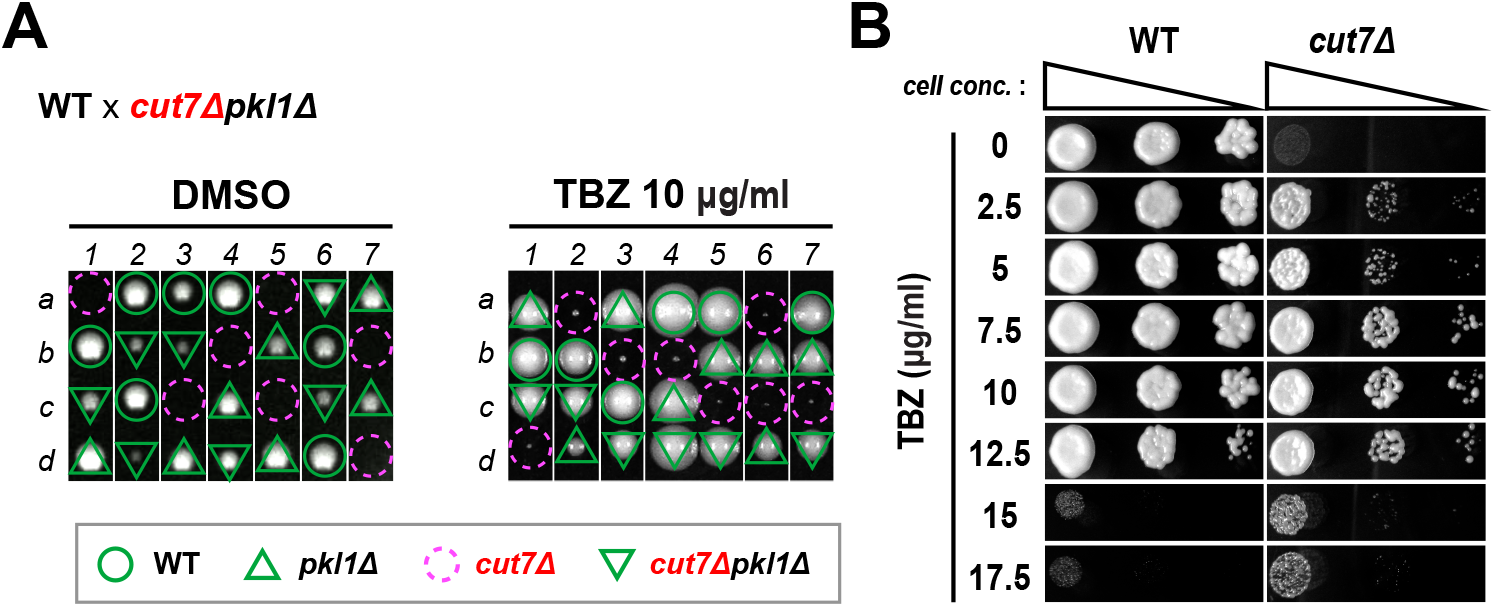
The microtubule-destabilizing drug bypasses the essential function of Cut7/Kinesin-5. **(A)** Tetrad analysis. Asci formed from diploid cells created by crossing between wild-type and *cut7Δpkl1Δ* strains were dissected on rich YE5S plates containing 1% DMSO (left) or 10 μg/ml TBZ (right), and spores were allowed to germinate and proliferate at 27°C for 4 d. Representative dissection patterns of individual spores (a–d) in each ascus (1–4) are shown. Assuming 2:2 segregation of individual markers allows the identification of genotype of each segregant. Circles in green indicate wild-type segregants, those in red show *cut7Δ,* triangles in green represent *pkl1Δ* and reversed triangles correspond to *cut7Δpkl1Δ.* Note that all *cut7Δ* spores formed viable, albeit small, colonies in the presence of TBZ, while they were inviable in its absence. **(B)** Spot test. One of wild-type or *cut7Δ* colonies obtained from tetrad dissection shown in **(A)** were spotted on YE5S plates in the absence or presence of various concentrations of TBZ, and incubated at 27°C for 3 d.

### Gene deletions perturbing spindle microtubule assembly are incapable of bypassing the essentiality of Cut7/Kinesin-5

Next, we addressed whether complete deletion of *skf* and other genes involved in microtubule assembly can also bypass the lethality caused by the *cut7* deletion. Tetrad dissection indicated that none of these deletions except for *pkl1* or *msd1* could rescue the lethality of *cut7Δ* cells (Figure 6A and B). As shown earlier (Figure 2), deletions of these genes led to the reduced levels of Klp2 on the spindles. Consistent with this, it is known that unlike *pkl1,* the *klp2* deletion is incapable of rescuing the lethality of *cut7Δ* cells (Yukawa *et al.* 2015). Note that unlike *pkl1* or *msd1, wdr8* deletion did not rescue the lethality, but *cut7Δwdr8Δklp2Δ* triple mutants were viable (data not shown). We envisage that even in the absence of Wdr8, cells retain residual Pkl1 activity, thereby rendering *cut7Δ* inviable only in the absence of Klp2. We, therefore, conclude that the reduction of Klp2 levels on spindle microtubules by TBZ treatment contributes to but is not sufficient for rescuing *cut7Δ* lethality.

**Figure 6.**
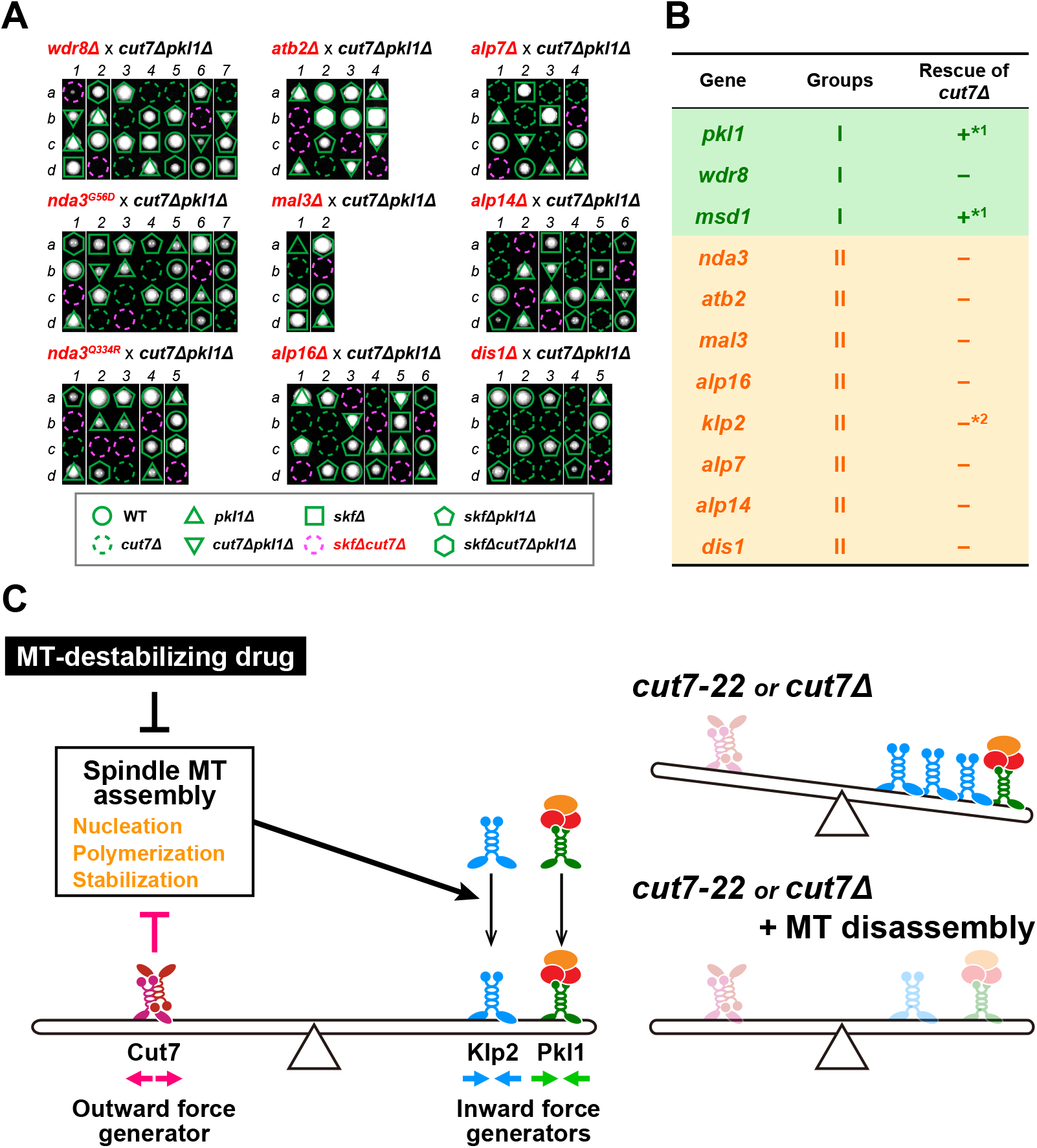
Inability of gene deletions to rescue *cut7-deleted* cells and a schematic model. **(A)** Tetrad analysis. Tetrad dissections upon crosses between *cut7Δpkl1Δ* and mutations or deletions of genes encoding tubulin or various MAPs (collectively shown as *skf* in an outlined frame at the bottom) were performed. Individual spores (a–d) in each ascus (1–4) were dissected on YE5S plates and incubated at 27°C for 3 d. Representative tetrad patterns are shown. Assuming 2:2 segregation of individual markers allows the identification of each segregant. Circles in green: wild type; Dotted circles in green: *cut7Δ;* Triangles in green: *pkl1Δ*; Reversed triangles in green: *cut7Δpkl1Δ*; Squares in green: *skfΔ;* Dotted circles in red: *skfΔcut7Δ;* Pentagons in green: *skfΔpkl1Δ*; Hexagons in green: *skfΔcut7Δpkl1Δ.* **(B)** A summary table of suppression profiles towards *cut7Δ.* *1: data from (Yukawa *et al.* 2017); *2: data from (Yukawa *et al.* 2018). **(C)** A model. Bipolar spindle formation requires collaborative balance between mitotic kinesins and microtubule stability/dynamics. Kinesin-5 (Cut7) generates outward force, while Kinesin-14s (Pkl1 and Klp2) generate opposing inward forces. Spindle microtubule stability/dynamics regulated by a cohort of MAPs play a positive role in Klp2 activity, while Cut7 plays an inhibitory role in microtubule nucleation, polymerization and/or stabilization, thereby suppressing Klp2 activity (left). In the *cut7-22* ts mutant, the levels of Klp2 on the mitotic spindles are upregulated. When *cut7-22* or *cut7Δ* cells are treated with a microtubule-depolymerizing drug, compromised microtubule dynamics and the reduced level of Klp2 on the spindle collaborate, by which these cells no longer require Cut7 for survival.

In summary, our genetic analysis indicates that two conditions can rescue hypomorphic *cut7* mutants. The first is the disruption of Pkl1/Kinesin-14 function, which is consistent with previous studies (Pidoux *et al.* 1996; Troxell *et al.* 2001; Olmsted *et al.* 2014; Syrovatkina and Tran 2015; Yukawa *et al.* 2017). The second is the impairment of spindle microtubules by mutations/deletions of genes encoding tubulins and a cohort of MAPs that are important for spindle assembly. We show that the second group lessens Klp2 localization on the mitotic spindle, which could account for the suppression of *cut7* ts mutants (Troxell *et al.* 2001; Yukawa *et al.* 2018). Markedly, continuous treatment by the microtubule-destabilizing drug TBZ is capable of not only suppressing the hypomorphic *cut7* mutant but also bypassing the essentiality of Cut7. As even complete deletion of *klp2* fails to rescue the *cut7* deletion (Yukawa *et al.* 2018), TBZ must interfere with some functions other than Klp2 downregulation. We posit that Klp2-independent pathway(s) regulating spindle microtubule stability/dynamics is also important to render Cut7 dispensable (Figure 6C).

## DISCUSSION

In this study, we have performed an unbiased genetic screening for suppressors of the *cut7-22* ts mutant, and uncovered a genetic network responsible for conferring lethality to this mutant. We have also found that treatment with the microtubule-depolymerizing drug TBZ rescues the *cut7-22* ts mutant. Remarkably, a low dose of TBZ is capable of even bypassing the essential function of Cut7. Hence, microtubule dynamics and/or assembly kinetics antagonize with Cut7, and the orchestration between these two factors is required for bipolar spindle assembly.

### Compromised microtubule stability/dynamics ameliorate Kinesin-5 deficiency

Previous work in HeLa cells has shown that the perturbation of microtubule stability/dynamics by either knockdown of the chTOG MAP (an ortholog of Alp14 and Dis1) or treatment with low concentrations of microtubule-disruptive drugs (Nocodazole and Vinblastin) can effectively rescue monopolar spindle phenotypes induced by Eg5/Kif11 inactivation (Kollu *et al.* 2009; Florian and Mayer 2011), and a similar observation is also reported in U2OS cells (Kollu *et al.* 2009). These results are wholly consistent with what we observe in fission yeast, indicating evolutionary conservation of Kinesin-5 function. It would, therefore, be worth investigating whether HSET/Kinesin-14 levels on the spindle microtubule are reduced under these conditions, which is observed for fission yeast Klp2/Kinesin-14 under analogous conditions.

### Cut7/Kinesin-5 regulates microtubule stability and/or dynamics

We show that in the *cut7-22* mutant, intensities of spindle microtubules are significantly increased. Furthermore, consistent with these data, *cut7-22* cells display hyper-resistance to microtubule-destabilizing TBZ (Yukawa *et al.* 2015). These results raise the intriguing question as to whether Kinesin-5 molecules per se play any role in microtubule stability and/or dynamics. Previous reports indeed show that Kinesin-5 regulates microtubule stability/dynamics as a microtubule depolymerizing factor (budding yeast Cin8 and human Eg5/Kif11) (Gardner *et al.* 2008; WANG *et al.* 2010), conversely as a microtubule stabilizer (budding yeast Kip1) (Fridman *et al.* 2013) or a microtubule polymerase *(Xenopus* Eg5) (Chen and Hancock 2015). In fission yeast, mutations in genes encoding the components of γ-TuC other than Alp16 are also capable of rescuing *cut7-22* (Rodriguez *et al.* 2008), and interestingly, Cut7/Kinesin-5 reportedly interacts with γ-TuC (Olmsted *et al.* 2014). It is, therefore, possible that by physically binding γ-TuC, Cut7 downregulates microtubule nucleating activity, which would explain the increased spindle intensities in the *cut7-22* mutant. As the the γ-TuC levels at the mitotic SPBs are unaltered in this mutant, Cut7 might act as a nucleation-inhibitory factor of the γ-TuC, instead of as a spatial regulator. It is noteworthy that it was previously claimed that Cut7 potentiates γ-TuC activity (Olmsted *et al.* 2014); however this notion is somewhat contradictory with genetic results obtained by the same authors (and this study), in which Cut7 and γ-TuC play opposing, rather than collaborative, roles.

Recently, it has become clear that the XMAP215/Dis1/TOG microtubule polymerase family is also a part of a microtubule nucleator; XMAP215 and Alp14 bind γ-TuC directly and indirectly, respectively, thereby promoting the very first step of microtubule assembly (Flor-Parra *et al.* 2018; Thawani *et al.* 2018). It is tempting to speculate that the microtubule nucleator complex includes both positive (the XMAP215/Dis1/TOG family) and negative regulators (Kinesin-5) in addition to its canonical γ-TuC components (Kollman *et al.* 2011; Paz and Luders 2018). Overall, whether and how Cut7 impacts γ-TuC-mediated microtubule nucleation and elongation is an outstanding issue that should be addressed in the future.

### A low dose of the microtubule-destabilizing reagent TBZ bypasses the requirement of Kinesin-5

One surprising observation made in this study is the rescue of *cut7Δ* cells by treatment with a low concentration of TBZ. As *cut7Δklp2Δ* double mutant cells are inviable, the inhibition of Klp2 activity with this drug treatment is not sufficient to fully account for the suppression, though it would make an important contribution. One possibility is the downregulation of Pkl1-mediated inward force. However, the measurement of Pkl1 intensities in TBZ-treated cells does not support this notion (Figure 4D). In both animal and fission yeast cells, treatment with low concentrations of microtubule-disruptive drugs interferes with microtubule dynamics and/or increases the concentration of free (unpolymerized) tubulins (Vasquez *et al.* 1997; Mary *et al.* 2015). Thus, compromised microtubule dynamics per se likely suppress the generation of lethal inward force in the absence of Kinesin-5, thereby rendering Kinesin-5 dispensable.

### Alp14 and Dis1 microtubule polymerases play both positive and negative roles in kinesin-dependent bipolar spindle assembly

We show here that the deletion of either *alp14* or *dis1* rescues the hypomorphic *cut7* mutant phenotype, indicating that Alp14/Dis1-dependent microtubule polymerization counteracts Cut7-mediated outward force. Curiously and apparently paradoxically, our previous work showed that either of these two MAPs becomes essential for cell viability in the absence of both Cut7 and Pkl1 *(cut7Δpkl1Δ)* (Yukawa *et al.* 2017), showing that Alp14 and Dis1 play a cooperative role with Cut7 in the absence of Kinesin-14. In this latter situation where Cut7 and Alp14/Dis1 (and other MAPs) functionally collaborate (Rincon *et al.* 2017; Yukawa *et al.* 2017), we previously argued that the plus ends of polymerizing microtubules push the respective opposite SPB, thereby generating outward force. This force is sufficient to separate two SPBs in *cut7Δpkl1Δ* cells.

In the hypomorphic *cut7* mutant cells, on the other hand, microtubule polymerization/stabilization mediated by Alp14 or Dis1 is deleterious to these cells. We hypothesize that this adverse impact elicited by Alp14 or Dis1 is mediated by Klp2, which generates antagonistic inward force against Cut7-dependent outward force (Yukawa *et al.* 2018). Taken together, bipolar spindle formation is under the control of a complex network consisting of multiple kinesins and MAPs, and individual molecules display different characters in a context-dependent manner.

### Concluding remarks

Much attention has been focused on mitotic kinesins as druggable targets in cancer therapeutics (Cosenza and Kramer 2016). In particular, inhibitors against Kinesins-5 and -14 are thought to be promising pharmacological reagents, and several small-molecule inhibitors have been clinically evaluated (Huszar *et al.* 2009). One of downsides, however, is the emergence of a cell population that is resistant to drug treatment. Results obtained in this work provide novel insight into the usage of these inhibitors; the combined treatment of Kinesin-5 inhibitors and microtubule stabilizing reagents such as Paclitaxel, which by itself is widely used in cancer chemotherapy (Shi *et al.* 2008; Mitchison 2012; Weaver 2014), would be beneficial for the treatment of drug resistant cancers.

## ACKNOWLEDGEMENTS

We thank Paul Nurse, Iain Hagan and J. Richard McIntosh for strains, and Tetsushi Iida and Masaru Ueno for the instruction for the isolation of genomic DNAs from revertants and data analysis of whole genome sequences with the Mudi method. We are grateful to Kylie Pan and Vismaya Kharkar for critical reading of the manuscript. This work was supported by the Japan Society for the Promotion of Science (JSPS) [KAKENHI Scientific Research (A) (16H02503 to T.T.), a Challenging Exploratory Research grant (16K14672 to T.T.), Scientific Research (C) (16K07694 to M.Y.)], the Naito Foundation (T.T.) and the Uehara Memorial Foundation (T.T).

## AUTHOR CONTRIBUTIONS

M.Y. and T.T. designed research. M.Y. and Y.Y. performed experiments and analyzed the data. M.Y. and T.T. wrote the manuscript, and Y.Y. made suggestions.

## REFERENCES

Al-Bassam, J., H. Kim, I. Flor-Parra, N. Lal, H. Velji et al., 2012 Fission yeast Alp14 is a dose dependent plus end tracking microtubule polymerase. Mol. Biol. Cell 23: 2878–2890.

Anders, A., P. C. Lourenco and K. E. Sawin, 2006 Noncore components of the fission yeast gamma-tubulin complex. Mol Biol Cell 17: 5075–5093.

Asakawa, K., M. Toya, M. Sato, M. Kanai, M. Kume et al., 2005 Mal3, the fission yeast EB1 homolog, cooperates with Bub1 spindle checkpoint to prevent monopolar attachment. EMBO Rep. 6: 1194–1120.

Bähler, J., J. Wu, M. S. Longtine, N. G. Shah, A. McKenzie III et al., 1998 Heterologous modules for efficient and versatile PCR-based gene targeting in *Schizosaccharomyces pombe*. Yeast 14: 943–951.

Beinhauer, J. D., I. M. Hagan, J. H. Hegemann and U. Fleig, 1997 Mal3, the fission yeast homologue of the human APC-interacting protein EB-1 is required for microtubule integrity and the maintenance of cell form. J Cell Biol 139: 717–728.

Blangy, A., H. A. Lane, P. d’Herin, M. Harper, M. Kress et al., 1995 Phosphorylation by p34^cdc2^ regulates spindle association of human Eg5, a kinesin-related motor essential for bipolar spindle formation in vivo. Cell 83: 1159–1169.

Braun, M., D. R. Drummond, R. A. Cross and A. D. McAinsh, 2009 The kinesin-14 Klp2 organizes microtubules into parallel bundles by an ATP-dependent sorting mechanism. Nat. Cell Biol. 11: 724–730.

Britto, M., A. Goulet, S. Rizvi, O. von Loeffelholz, C. A. Moores et al., 2016 *Schizosaccharomyces pombe* kinesin-5 switches direction using a steric blocking mechanism. Proc Natl Acad Sci U S A 113: E7483–E7489.

Carvalho, P., J. S. Tirnauer and D. Pellman, 2003 Surfing on microtubule ends. Trends Cell Biol. 13: 229–237.

Chen, Y., and W. O. Hancock, 2015 Kinesin-5 is a microtubule polymerase. Nat Commun 6: 8160.

Cosenza, M. R., and A. Kramer, 2016 Centrosome amplification, chromosomal instability and cancer: mechanistic, clinical and therapeutic issues. Chromosome Res 24: 105–126.

Dumas, M. E., E. G. Sturgill and R. Ohi, 2016 Resistance is not futile: Surviving Eg5 inhibition. Cell Cycle 15: 2845–2847.

Edamatsu, M., 2014 Bidirectional motility of the fission yeast kinesin-5, Cut7. Biochem. Biophys. Res. Commun. 446: 231–234.

Enos, A. P., and N. R. Morris, 1990 Mutation of a gene that encodes a kinesin-like protein blocks nuclear division in *A. nidulans*. Cell 60: 1019–1027.

Flor-Parra, I., A. B. Iglesias-Romero and F. Chang, 2018 The XMAP215 ortholog Alp14 promotes microtubule nucleation in fission yeast. Curr Biol 28: 1681–1691.e1684.

Florian, S., and T. U. Mayer, 2011 Modulated microtubule dynamics enable Hklp2/Kif15 to assemble bipolar spindles. Cell Cycle 10: 3533–3544.

Fridman, V., A. Gerson-Gurwitz, O. Shapira, N. Movshovich, S. Lakamper et al., 2013 Kinesin-5 Kip1 is a bi-directional motor that stabilizes microtubules and tracks their plus-ends in vivo. J Cell Sci. 126: 4147–4159.

Fu, C., J. J. Ward, I. Loiodice, G. Velve-Casquillas, F. J. Nedelec et al., 2009 Phospho-regulated interaction between kinesin-6 Klp9p and microtubule bundler Ase1p promotes spindle elongation. Dev. Cell 17: 257–267.

Fujita, A., L. Vardy, M. A. Garcia and T. Toda, 2002 A fourth component of the fission yeast γ-tubulin complex, Alp16, is required for cytoplasmic microtubule integrity and becomes indispensable when γ-tubulin function is compromised. Mol. Biol. Cell 13: 2360–2373.

Furuta, K., M. Edamatsu, Y. Maeda and Y. Y. Toyoshima, 2008 Diffusion and directed movement: In vitro motile properties of fission yeast kinesin-14 Pkl1. J. Biol. Chem. 283: 36465–36473.

Garcia, M. A., L. Vardy, N. Koonrugsa and T. Toda, 2001 Fission yeast ch-TOG/XMAP215 homologue Alp14 connects mitotic spindles with the kinetochore and is a component of the Mad2-dependent spindle checkpoint. EMBO J. 20: 3389–3401.

Gardner, M. K., D. C. Bouck, L. V. Paliulis, J. B. Meehl, E. T. O’Toole et al., 2008 Chromosome congression by Kinesin-5 motor-mediated disassembly of longer kinetochore microtubules. Cell 135: 894–906.

Gerson-Gurwitz, A., C. Thiede, N. Movshovich, V. Fridman, M. Podolskaya et al., 2011 Directionality of individual kinesin-5 Cin8 motors is modulated by loop 8, ionic strength and microtubule geometry. EMBO J. 30: 4942–4954.

Hagan, I., and M. Yanagida, 1990 Novel potential mitotic motor protein encoded by the fission yeast *cut7^+^* gene. Nature 347: 563–566.

Hagan, I., and M. Yanagida, 1992 Kinesin-related cut7 protein associates with mitotic and meiotic spindles in fission yeast. Nature 356: 74–76.

Heck, M. M., A. Pereira, P. Pesavento, Y. Yannoni, A. C. Spradling et al., 1993 The kinesin-like protein KLP61F is essential for mitosis in *Drosophila*. J Cell Biol 123: 665–679.

Hiraoka, Y., T. Toda and M. Yanagida, 1984 The *NDA3* gene of fission yeast encodes β-tubulin: a cold-sensitive *nda3* mutation reversibly blocks spindle formation and chromosome movement in mitosis. Cell 39: 349–358.

Hussmann, F., D. R. Drummond, D. R. Peet, D. S. Martin and R. A. Cross, 2016 Alp7/TACC-Alp14/TOG generates long-lived, fast-growing MTs by an unconventional mechanism. Sci Rep 6: 20653.

Huszar, D., M. E. Theoclitou, J. Skolnik and R. Herbst, 2009 Kinesin motor proteins as targets for cancer therapy. Cancer Metastasis Rev 28: 197–208.

Iida, N., F. Yamao, Y. Nakamura and T. Iida, 2014 Mudi, a web tool for identifying mutations by bioinformatics analysis of whole-genome sequence. Genes Cells 19: 517–527.

Ikebe, C., M. Konishi, D. Hirata, T. Matsusaka and T. Toda, 2011 Systematic localization study on novel proteins encoded by meiotically up-regulated ORFs in fission yeast. Biosci. Biotechnol. Biochem. 75: 2364–2370.

Kapitein, L. C., E. J. Peterman, B. H. Kwok, J. H. Kim, T. M. Kapoor et al., 2005 The bipolar mitotic kinesin Eg5 moves on both microtubules that it crosslinks. Nature 435: 114–118.

Kapoor, T. M., T. U. Mayer, M. L. Coughlin and T. J. Mitchison, 2000 Probing spindle assembly mechanisms with monastrol, a small molecule inhibitor of the mitotic kinesin, Eg5. J. Cell Biol. 150: 975–988.

Kashina, A. S., R. J. Baskin, D. G. Cole, K. P. Wedaman, W. M. Saxton et al., 1996 A bipolar kinesin. Nature 379: 270–272.

Kollman, J. M., A. Merdes, L. Mourey and D. A. Agard, 2011 Microtubule nucleation by γ-tubulin complexes. Nat. Rev. Mol. Cell Biol. 12: 709–721.

Kollu, S., S. F. Bakhoum and D. A. Compton, 2009 Interplay of microtubule dynamics and sliding during bipolar spindle formation in mammalian cells. Curr Biol 19: 2108–2113.

Le Guellec, R., J. Paris, A. Couturier, C. Roghi and M. Philippe, 1991 Cloning by differential screening of a *Xenopus* cDNA that encodes a kinesin-related protein. Mol Cell Biol 11: 3395–3398.

Loiodice, I., J. Staub, T. Gangi-Setty, N. P. Nguyen, A. Paoletti et al., 2005 Ase1p organizes anti-parallel microtubule arrays during interphase and mitosis in fission yeast. Mol. Biol. Cell 16: 1756–1768.

Ma, H. T., S. Erdal, S. Huang and R. Y. Poon, 2014 Synergism between inhibitors of Aurora A and KIF11 overcomes KIF15-dependent drug resistance. Mol Oncol 8: 1404–1418.

Mana-Capelli, S., J. R. McLean, C. T. Chen, K. L. Gould and D. McCollum, 2012 The kinesin-14 Klp2 is negatively regulated by the SIN for proper spindle elongation and telophase nuclear positioning. Mol. Biol. Cell 23: 4592–4600.

Mary, H., J. Fouchard, G. Gay, C. Reyes, T. Gauthier et al., 2015 Fission yeast kinesin-8 controls chromosome congression independently of oscillations. J Cell Sci 128: 3720–3730.

Masuda, H., and T. Toda, 2016 Synergistic role of fission yeast Alp16^GCP6^ and Mzt1^MOZART1^ in γ-tubulin complex recruitment to mitotic spindle pole bodies and spindle assembly. Mol Biol Cell 27: 1753–1763.

Matsuo, Y., S. P. Maurer, M. Yukawa, S. Zakian, M. R. Singleton et al., 2016 An unconventional interaction between Dis1/TOG and Mal3/EB1 in fission yeast promotes the fidelity of chromosome segregation. J Cell Sci 129: 4592–4606.

Maundrell, K., 1990 *nmt1* of fission yeast. A highly transcribed gene completely repressed by thiamine. J Biol Chem 265: 10857–10864.

Mitchison, T. J., 2012 The proliferation rate paradox in antimitotic chemotherapy. Mol. Biol. Cell 23: 1–6.

Mitchison, T. J., and E. D. Salmon, 2001 Mitosis: a history of division. Nat Cell Biol 3: E17–E21.

Moreno, S., A. Klar and P. Nurse, 1991 Molecular genetic analysis of fission yeast *Schizosaccharomyces pombe*. Methods Enzymol 194: 795–823.

Nabeshima, K., H. Kurooka, M. Takeuchi, K. Kinoshita, Y. Nakaseko et al., 1995 p93^dis1^, which is required for sister chromatid separation, is a novel microtubule and spindle pole body-associating protein phosphorylated at the Cdc2 target sites. Genes Dev. 9: 1572–1585.

Olmsted, Z. T., A. G. Colliver, T. D. Riehlman and J. L. Paluh, 2014 Kinesin-14 and kinesin-5 antagonistically regulate microtubule nucleation by γ-TuRC in yeast and human cells. Nat Commun 5: 5339.

Paz, J., and J. Luders, 2018 Microtubule-organizing centers: towards a minimal parts list. Trends Cell Biol 28: 176–187.

Pidoux, A. L., M. LeDizet and W. Z. Cande, 1996 Fission yeast pkl1 is a kinesis-related protein involved in mitotic spindle function. Mol. Biol. Cell 7: 1639–1655.

Rincon, S. A., A. Lamson, R. Blackwell, V. Syrovatkina, V. Fraisier et al., 2017 Kinesin-5-independent mitotic spindle assembly requires the antiparallel microtubule crosslinker Ase1 in fission yeast. Nat Commun 8: 15286.

Rodriguez, A. S., J. Batac, A. N. Killilea, J. Filopei, D. R. Simeonov et al., 2008 Protein complexes at the microtubule organizing center regulate bipolar spindle assembly. Cell Cycle 7: 1246–1253.

Roostalu, J., C. Hentrich, P. Bieling, I. A. Telley, E. Schiebel et al., 2011 Directional switching of the kinesin Cin8 through motor coupling. Science 332: 94–99.

Sato, M., S. Dhut and T. Toda, 2005 New drug-resistant cassettes for gene disruption and epitope tagging in *Schizosaccharomyces pombe*. Yeast 22: 583–591.

Sato, M., L. Vardy, M. A. Garcia, N. Koonrugsa and T. Toda, 2004 Interdependency of fission yeast Alp14/TOG and coiled coil protein Alp7 in microtubule localization and bipolar spindle formation. Mol. Biol. Cell 15: 1609–1622.

Sawin, K. E., K. LeGuellec, M. Philippe and T. J. Mitchison, 1992 Mitotic spindle organization by a plus-end-directed microtubule motor. Nature 359: 540–543.

Shapira, O., A. Goldstein, J. Al-Bassam and L. Gheber, 2017 A potential physiological role for bi-directional motility and motor clustering of mitotic kinesin-5 Cin8 in yeast mitosis. J Cell Sci 130: 725–734.

She, Z. Y., and W. X. Yang, 2017 Molecular mechanisms of kinesin-14 motors in spindle assembly and chromosome segregation. J Cell Sci 130: 2097–2110.

Shi, J., J. D. Orth and T. Mitchison, 2008 Cell type variation in responses to antimitotic drugs that target microtubules and kinesin-5. Cancer Res. 68: 3269–3276.

Syrovatkina, V., and P. T. Tran, 2015 Loss of kinesin-14 results in aneuploidy via kinesin-5-dependent microtubule protrusions leading to chromosome cut. Nat Commun 6: 7322.

Thawani, A., R. S. Kadzik and S. Petry, 2018 XMAP215 is a microtubule nucleation factor that functions synergistically with the gamma-tubulin ring complex. Nat Cell Biol 20: 575–585.

Toda, T., Y. Adachi, Y. Hiraoka and M. Yanagida, 1984 Identification of the pleiotropic cell division cycle gene *NDA2* as one of two different α-tubulin genes in *Schizosaccharomyces pombe*. Cell 37: 233–242.

Toya, M., M. Sato, U. Haselmann, K. Asakawa, D. Brunner et al., 2007 γ-Tubulin complex-mediated anchoring of spindle microtubules to spindle-pole bodies requires Msd1 in fission yeast. Nat Cell Biol 9: 646–653.

Troxell, C. L., M. A. Sweezy, R. R. West, K. D. Reed, B. D. Carson et al., 2001 *pkl1+* and *klp2*^+^: two kinesins of the Kar3 subfamily in fission yeast perform different functions in both mitosis and meiosis. Mol. Biol. Cell 12: 3476–3488.

Vardy, L., and T. Toda, 2000 The fission yeast γ-tubulin complex is required in G_1_ phase and is a component of the spindle assembly checkpoint. EMBO J. 19: 6098–6111.

Vasquez, R. J., B. Howell, A. M. Yvon, P. Wadsworth and L. Cassimeris, 1997 Nanomolar concentrations of nocodazole alter microtubule dynamic instability in vivo and in vitro. Mol Biol Cell 8: 973–985.

Wacker, S. A., B. R. Houghtaling, O. Elemento and T. M. Kapoor, 2012 Using transcriptome sequencing to identify mechanisms of drug action and resistance. Nat Chem Biol 8: 235–237.

Wang, G., X. Gao, Y. Huang, Z. Yao, Q. Shi et al., 2010 NPM/B23 inhibits Eg5-mediated microtubules depolymerization via inactivating its ATPase activity. J. Biol. Chem. 285: 19060–19067.

Weaver, B. A., 2014 How Taxol/paclitaxel kills cancer cells. Mol. Biol. Cell 25: 2677–2681.

West, R. R., E. V. Vaisberg, R. Ding, P. Nurse and J. R. McIntosh, 1998 *cut11+:* a gene required for cell cycle-dependent spindle pole body anchoring in the nuclear envelope and bipolar spindle formation in *Schizosaccharomyces pombe*. Mol. Biol. Cell 9: 2839–2855.

Woodruff, J. B., O. Wueseke and A. A. Hyman, 2014 Pericentriolar material structure and dynamics. Philos. Trans. R. Soc. Lond. B. Biol. Sci. 369.

Yamashita, A., M. Sato, A. Fujita, M. Yamamoto and T. Toda, 2005 The roles of fission yeast ase1 in mitotic cell division, meiotic nuclear oscillation, and cytokinesis checkpoint signaling. Mol Biol Cell 16: 1378–1395.

Yount, A. L., H. Zong and C. E. Walczak, 2015 Regulatory mechanisms that control mitotic kinesins. Exp Cell Res 334: 70–77.

Yukawa, M., C. Ikebe and T. Toda, 2015 The Msd1-Wdr8-Pkl1 complex anchors microtubule minus ends to fission yeast spindle pole bodies. J Cell Biol 209: 549–562.

Yukawa, M., T. Kawakami, M. Okazaki, K. Kume, N. H. Tang et al., 2017 A microtubule polymerase cooperates with the kinesin-6 motor and a microtubule cross-linker to promote bipolar spindle assembly in the absence of kinesin-5 and kinesin-14 in fission yeast. Mol Biol Cell 28: 3647–3659.

Yukawa, M., Y. Yamada, T. Yamauchi and T. Toda, 2018 Two spatially distinct kinesin-14 proteins, Pkl1 and Klp2, generate collaborative inward forces against kinesin-5 Cut7 in *S. pombe*. J Cell Sci 131: 1–11.

